# Estimating statistical power for event-related potential studies using the late positive potential

**DOI:** 10.1101/574368

**Authors:** Kyla D. Gibney, George Kypriotakis, Paul M. Cinciripini, Jason D. Robinson, Jennifer A. Minnix, Francesco Versace

## Abstract

The late positive potential (LPP) is a common measurement used to study emotional processes of subjects in event-related potential (ERP) paradigms. Despite its extensive use in affective neuroscience, there is presently no gold standard for how to appropriately power ERP studies using the LPP in within-subject and between-subjects experimental designs. The present study investigates how the number of trials, number of subjects, and magnitude of the effect size affect statistical power in analyses of the LPP. Using Monte Carlo simulations of ERP experiments with varying numbers of trials, subjects, and effect sizes, we measured the probability of obtaining a statistically significant effect in 1,489 different experiments repeated 1,000 times each. Predictably, our results showed that statistical power increases with increasing numbers of trials and subjects and at larger effect sizes. In addition, we found that higher levels of statistical power can be achieved with lower numbers of subjects and trials and at lower effect sizes in within-subject than in between-subjects designs. Furthermore, we found that, as subjects are added to an experiment, the slope of the relationship between effect size and statistical power increases and shifts to the left until the power asymptotes to nearly 100% at higher effect sizes. This suggests that adding more subjects greatly increases statistical power at lower effect sizes (<1 µV) compared with more robust (>1.5 µV) effect sizes.

## 1. Introduction

Identifying emotions’ neural correlates has high clinical relevance because most psychological disorders are characterized by altered affective processes (American Psychiatric Association, 2013; Dunning et al., 2011; Moeller et al., 2014; Oliver, Jentink, Drobes, & Evans, 2016; Simmons et al., 2008; Wirkner et al., 2017; X. Zhang et al., 2014), and associating neural responses to specific symptoms and behaviors is likely to inform and foster the development of new evidence-based treatments (Dunning et al., 2011; Robbins, Gillan, Smith, de Wit, & Ersche, 2012; Wirkner et al., 2017).

One of the most commonly used experimental paradigms for studying affective processes is the picture-viewing paradigm, sometimes described as the cue-reactivity paradigm (Uhrig et al., 2016). In its basic implementation, the paradigm requires participants to look at pleasant, unpleasant, and neutral images while their responses to these images are recorded. Thanks to the availability of several sets of standardized pictures (e.g., the International Affective Picture System, Bradley, M.M. & Lang, P.J. 2007; the Open Libray of Affective Foods, Miccoli et al., 2014; and EmoPicS, Wessa, Neumeister, Kanske, & Schonfelder, 2010), scientists can assess affective responses to a multitude of stimuli using self-reports (Bradley, M.M. & Lang, P.J.1994), peripheral (M. M. Bradley, Codispoti, Cuthbert, & Lang, 2001) and central (Peter J. Lang & Bradley, 2010) measures of nervous system activity.

When researchers record the event-related potentials (ERPs) elicited by emotional images, they often focus their analyses on the amplitude of the late positive potential (LPP) (Olofsson, Nordin, Sequeira, & Polich, 2008). The LPP, a sustained ERP component elicited by emotionally relevant images over central and parietal electrode sites, peaks between 400 and 800 ms after picture onset (Codispoti, Mazzetti, & Bradley, 2009; Cuthbert, Schupp, Bradley, Birbaumer, & Lang, 2000; Schupp et al., 2004; Versace et al., 2011). Both pleasant and unpleasant stimuli increase the amplitude of the LPP as a function of emotional arousal (Minnix et al., 2013; Weinberg & Hajcak, 2010). The affective modulation of the LPP is not influenced by the images’ perceptual composition (MM Bradley, Löw, & Lang, 2007; Codispoti, De Cesarei, & Ferrari, 2012; De Cesarei & Codispoti, 2006), exposure time (Codispoti et al., 2009), or repetition (Ferrari, Codispoti, & Bradley, 2017). These characteristics lead researchers to consider the LPP a robust index of motivational relevance (Peter J. Lang & Bradley, 2010) and a useful tool for investigating affective processes in clinical populations, such as individuals affected by neurological disorders (Xie et al., 2018), PTSD and other anxiety disorders (Fitzgerald et al., 2018; MacNamara, Jackson, Fitzgerald, Hajcak, & Phan, 2018; B.-W. Zhang et al., 2016), depression (Weinberg, Perlman, Kotov, & Hajcak, 2016), drug abuse (Dunning et al., 2011; Versace et al., 2017), anhedonia (Weinberg et al., 2016), psychopathy (Ellis, Schroder, Patrick, & Moser, 2017), and schizophrenia (Culbreth, Foti, Barch, Hajcak, & Kotov, 2018; Strandburg et al., 1994).

Researchers studying affective processes may use within-subject designs to evaluate treatment effects, or they may use between-subjects designs to compare affective responses across populations. However, designing these studies requires careful examination of statistical power, which has not been extensively examined in the current literature. ERP researchers may elect designs with small sample sizes or reduced number of trials in order to maximize recruitment while reducing subject burden. These decisions have an uncertain impact on statistical power and may reduce the likelihood of drawing appropriate conclusions from the experiment. As Ioannidis and others have stated, it is likely that the conclusions of many scientific studies are in fact false because of the studies’ insufficient power (Ioannidis, 2005; Button et al., 2013; Munafò et al., 2017). Hence, without systematically investigating how factors such as the number of subjects, the number of trials per condition, and the effect size investigated affect the statistical power of experiments that use the LPP as a dependent variable, it is difficult to draw reliable conclusions that can be translated to clinical applications in the psychological domain.

Presently there is no gold standard to estimate the statistical power of studies using the LPP in within-subject or between-subjects experimental designs. Researchers often have questions, such as: How many subjects per group are necessary to reliably determine whether two groups differ in their emotional responses? How many trials per condition are necessary to reliably determine whether two categories of stimuli evoke LPPs of different amplitudes? And given that different stimulus categories can evoke LPPs with very different amplitudes, how do the numbers of trials and subjects affect statistical power as a function of effect size? To address these questions, we adapted the simulation approach used by Boudewyn and colleagues to investigate statistical power for the error-related negativity (ERN) and the lateralized readiness potential (LRP) (Boudewyn, Luck, Farrens, & Kappenman, 2018). Briefly, Boudewyn and colleagues conducted Monte Carlo simulations of ERP experiments using real electroencephalogram data. Drawing from a main dataset, they simulated ERP experiments by randomly sampling subsets with varying numbers of trials, varying numbers of subjects, and varying effect sizes. Each simulated experiment was repeated 1,000 times and then tested for statistical significance. This procedure allowed the authors to calculate the probability of obtaining a statistically significant result at a significance level of 0.05 (i.e., the statistical power) for each combination of parameters.

While informative, the results that Boudewyn et al. obtained for the ERN and the LRP cannot be extrapolated to the LPP, as these components have different characteristics. For example, while the LPP is a stimulus-locked component elicited by emotionally relevant images, the ERN is a response-locked component elicited by the subject’s mistakes made during a behavioral task, and the LRP is a component associated with the preparation of a manual response (Luck, 2014). Furthermore, Boudewyn and colleagues studied undergraduate college students using active electrodes; these conditions cannot always be replicated in a clinical laboratory with older, often less healthy community participants. Because of issues such as patient comfort, sponge-based electrode systems are often favored in clinical environments, as they facilitate the electrode application process and eliminate the need to use gel, which some participants find uncomfortable. However, sponge-based systems are subject to a greater degree of noise than systems using traditional or active electrodes (Jackson & Bolger, 2014; Kappenman & Luck, 2010), so their use involves sacrificing some data quality in favor of patient comfort. Given the less-than-ideal recording conditions that some clinical laboratories might experience, using an appropriate dataset to evaluate the tradeoffs between the number of subjects, number of trials, and reliability of the results is highly relevant.

In this study, we conducted our simulations on LPP data collected from a large sample (N=314) of community participants with a wide age range (18 to 65 years), and we used a sponge-based electrode system coupled with a high impedance amplifier. As such, the results presented here should assist affective neuroscientists in maintaining the critical balance between patient comfort, rigor, and reproducibility and thereby further the development of evidence-based interventions in psychological disorders.

## 2. Methods

### 2.1. Study participants

For the analyses presented here, we used data from 314 community participants whom we had previously recruited for clinical studies of smoking cessation (Stevens et al., 2019). The data included here were recorded at baseline, before the beginning of any treatment. All participants were recruited from the Houston metro area through newspaper and radio advertisements. Inclusion criteria for the studies were: age, 18-65 years; fluent English speaker; not taking psychotropic medication; not diagnosed with a psychiatric disorder; and not having an uncontrolled medical illness. One subject was later excluded from the analyses because of incomplete data, leaving 313 subjects who were included in our analysis. Demographic information for our subjects is provided in Table 1. Each subject provided informed consent, and the study was approved by The University of Texas MD Anderson Cancer Center’s Institutional Review Board.

**Table 1:**
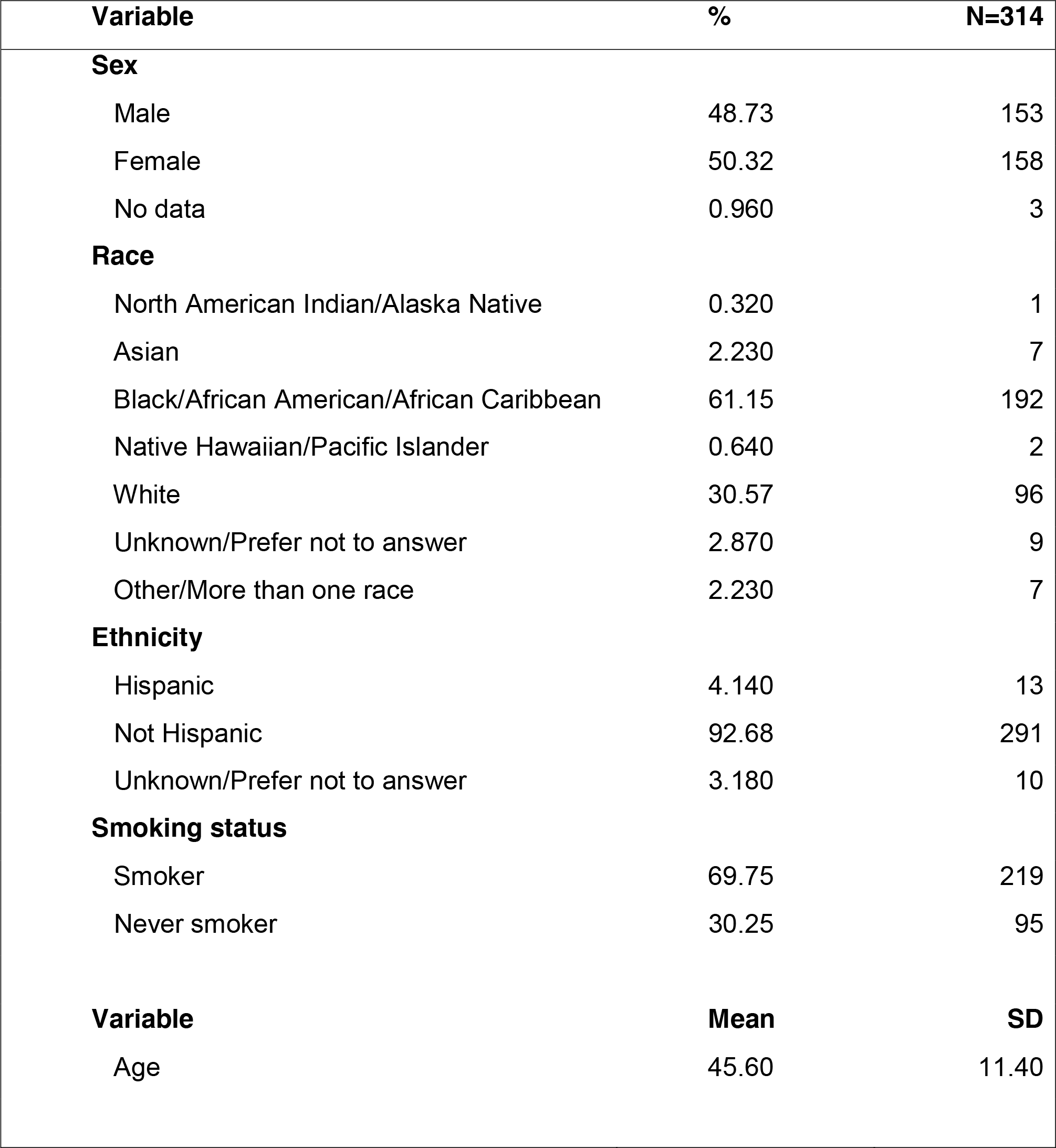
Baseline demographic information.

### 2.2. Picture-viewing task

The picture-viewing task used in this study included 192 images selected from the International Affective Picture System (P.J. Lang, Bradley, & Cuthbert, 2008) and from other picture collections previously used by our lab (Carter et al., 2006; Versace et al., 2011). The set included four picture categories: pleasant, unpleasant, cigarette-related, and neutral. Each category included 48 images (pleasant: 16 erotic scenes, 16 romantic couples, and 16 food images; unpleasant: 16 showing mutilations, 16 showing violence, and 16 showing accidents; cigarette-related: 32 of people smoking and 16 of smoking paraphernalia; neutral: 32 of people engaged in mundane activities and 16 household objects). During the task, images were presented in pseudo-random orders with no more than two pictures of the same category presented consecutively. Each picture was presented for 4 s, constituting a trial, and was followed by an inter-trial interval varying between 3 and 5 s, during which the subjects saw a black background with a white fixation cross. The entire picture presentation and recording session lasted approximately 30 min. Sessions were divided into three 10-min blocks separated by a 30-s break between blocks. Stimuli were presented using E-Prime 1 (PST Inc., Pittsburgh, PA) stimulus presentation software (Schneider, Eschman, & Zuccolotto, 2002) on a 42” plasma screen placed approximately 1.5 m from the participants’ eyes. Images were subtended horizontally at a horizontal visual angle of approximately 24 degrees.

### 2.3. Data collection procedures

During the picture presentation, ERPs were recorded using a 129-channel geodesic sensor net amplified with an AC-coupled 200-MΩ impedance amplifier (EGI Geodesic EEG System 200; Electrical Geodesics, Inc., Eugene, OR, USA) and referenced to Cz. Data were sampled at a rate of 250 Hz and were filtered online using a 0.1-Hz high-pass filter and a 100-Hz low-pass filter. As per the manufacturer’s instructions, scalp impedance was below 50 KΩ at the beginning of the recording.

### 2.4. Data reduction procedures

Even though we used only data from neutral trials in the Monte Carlo simulations (see Section 2.6), we reduced the data and plotted the results of both emotional and neutral categories to ensure that the data used in our analyses belonged to a standard LPP experiment. First, we corrected eye blink artifacts using a spatial filtering method as implemented in BESA software (BESA GmbH, Gräfelfing, Germany) and transformed the data to the average reference. Then, we imported the data into Brain Vision Analyzer 2.1 (Brain Products GmbH, Gilching, Germany) and filtered them with a low-pass filter of 30 Hz, a high-pass filter of 0.1 Hz, and a notch filter of 60 Hz. The data were then segmented into 900-ms segments, starting 100 ms before stimulus presentation. The 100-ms interval before stimulus presentation was defined as the baseline and subtracted from every data point in the segments. Artifacts were identified in the segmented data and were defined by 1) an amplitude of above 100 µV or below - 100 µV; 2) an absolute difference of greater than 100 µV between any two data points in a single segment; and 3) a maximum gradient of 25 µV/ms voltage step. Channels contaminated by artifacts in more than 40% of the segments were interpolated using six neighboring channels. We averaged the voltage from 10 centroparietal sensors (EGI electrodes 7, 31, 37, 54, 55, 79, 80, 87, 106, 129) because, in previous studies, these channels had shown the highest LPP differences between experimental conditions (Versace et al., 2011). We checked for the presence of artifacts in the averaged data using the same criteria mentioned above and discarded the segments contaminated by artifacts. Then, data from subjects with less than 40 artifact-free, neutral trials were removed, leaving data from 313 subjects with no artifacts. For each of the 313 subjects, we calculated a grand mean ERP and 95% confidence interval for each picture category (Figure 1). We calculated the LPP as the average voltages between 400 and 800 ms after stimulus onset for each picture subcategory within the pooled sensors. As expected, images with high motivational relevance, such as erotic or mutilation images, prompted higher LPPs than images with low motivational relevance, such as household objects (Figure 2).

**Figure 1:**
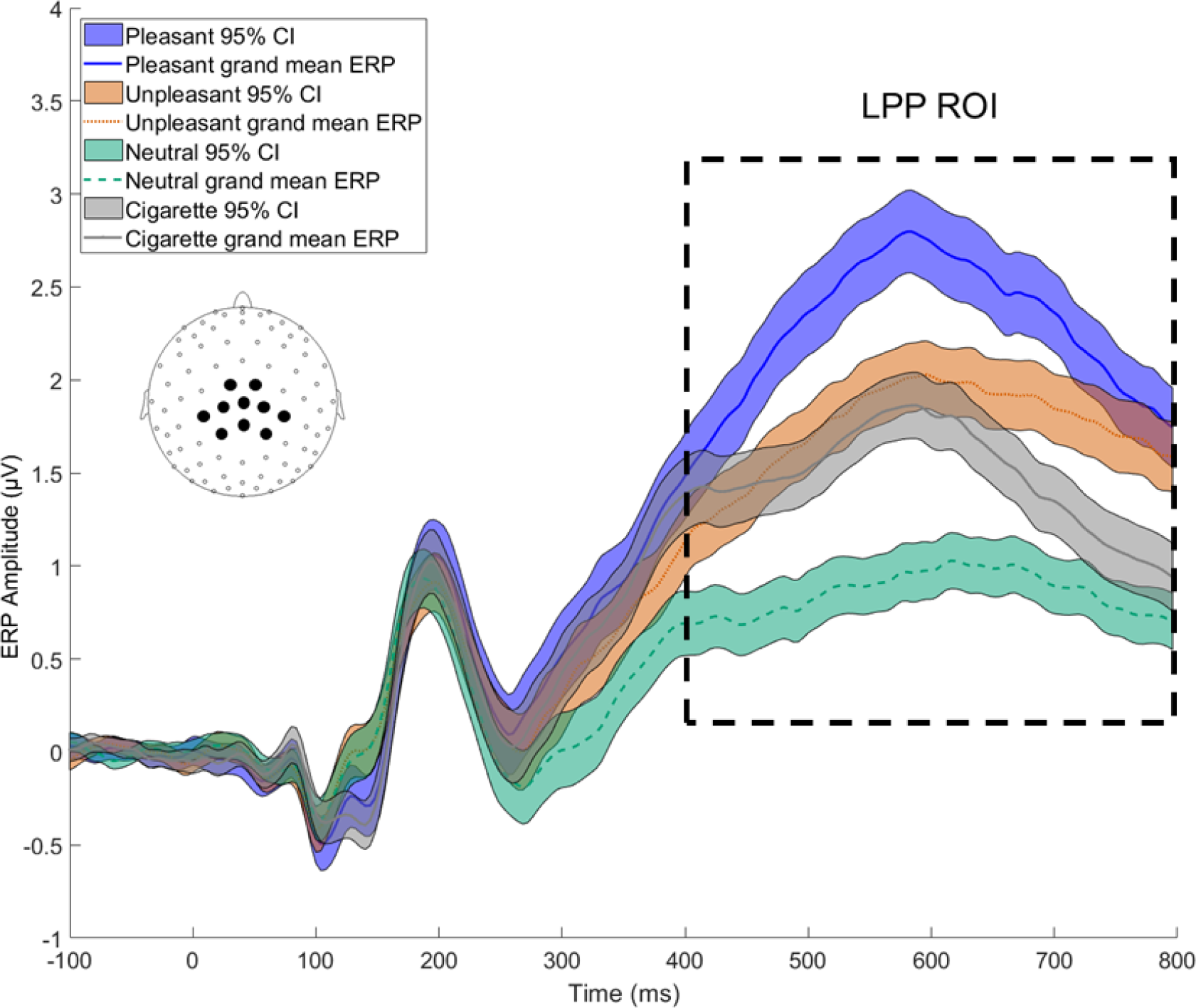
Grand mean ERPs (colored lines) and 95% confidence intervals (shaded) by category. The ERPs evoked by each category are consistent with previous studies, as more emotionally relevant picture categories evoke larger LPPs. CI, confidence interval; ERP, event-related potential; LPP, late positive potential; ROI, region of interest.

**Figure 2:**
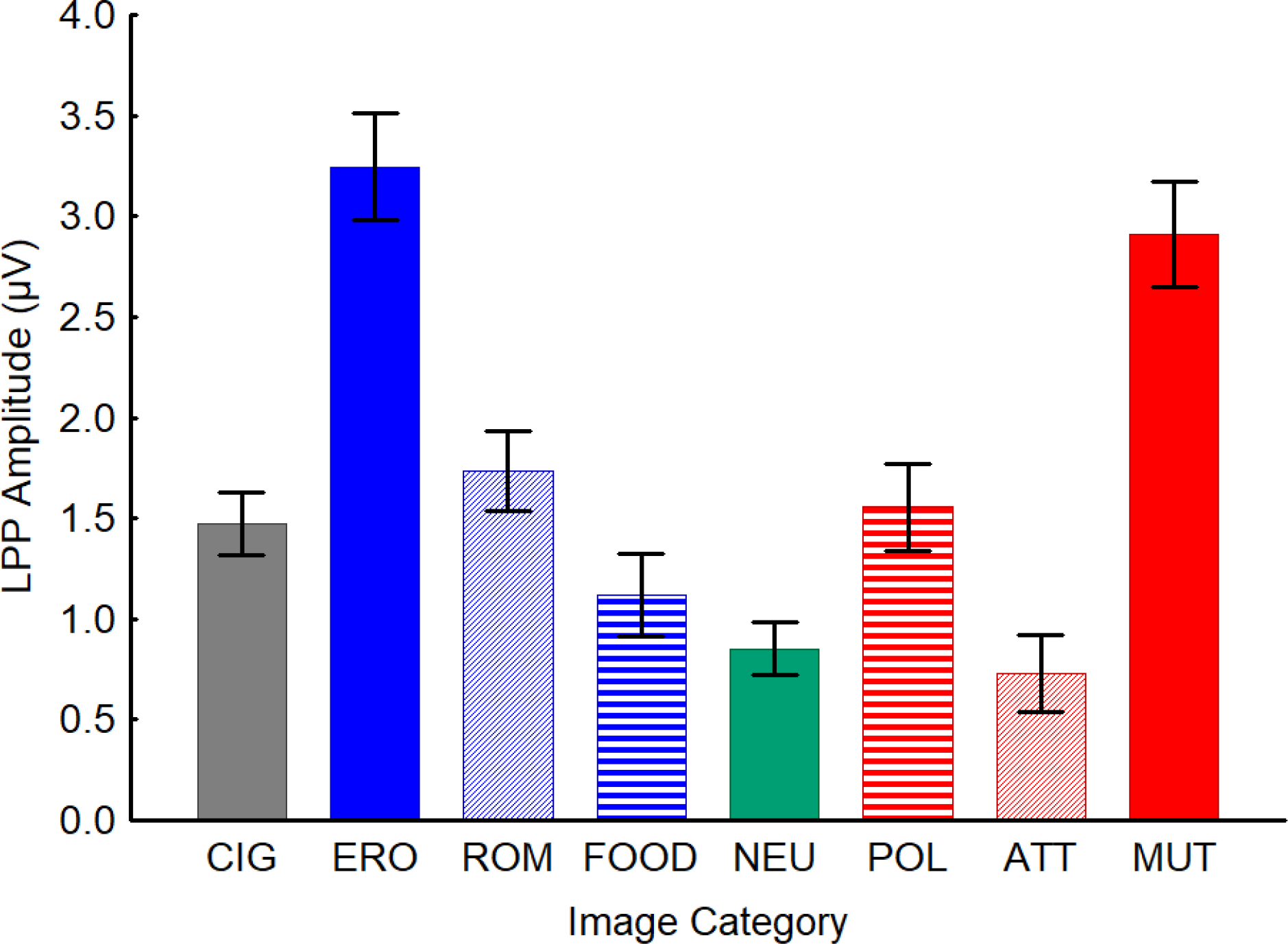
Average LPPs by category for all subjects (N= 313). The LPPs evoked by our stimuli were consistent with previous findings regarding the LPP: Emotionally relevant images evoke greater LPPs than less salient images. LPP, late positive potential. ERO, erotica. ROM, romance. FOOD, food. NEU, neutral. POL, pollution. VIOL, violence. MUT, mutilations. CIG, cigarettes.

### 2.5. Noise visualization

Before proceeding with the Monte Carlo simulations, we assessed the level of noise in the neutral trials. To visualize the noise in the segmented data, we followed the plus-minus averaging procedure outlined by Boudewyn et al. (Boudewyn et al., 2018; Schimmel, 1967). The goal of the procedure is to cancel the ERP signal and leave only the noise in the data. First, for each subject, we separated the time series data from all odd and even neutral trials into unique vectors and averaged them individually. Then, for each subject, we subtracted the mean of the odd trials from each even trial and vice versa. We tested the success of the procedure by averaging the time series data for all the trials (N = 11,328) after the subtraction. The average (shown in Figure 3) ranged from 1.5×10^−15^ to −1.5×10^−15^ µV, and thus remained at approximately zero, indicating that all the signal had been subtracted from the data, leaving only noise in the traces (Figure 3). To provide a more readable quantitative estimation of the variability around the mean, we calculated the percentage of trials at each time point that fell into each of four voltage bins: ±1 µV, ±5 µV, ±10 µV, and ±20 µV (Figure 4). At any given point, approximately 98% of the trials fell into the ± 20-µV range, leaving 2% beyond that range. We decided to keep these outliers in the analysis to model the degree of noise typical in data collected in a clinical setting.

**Figure 3:**
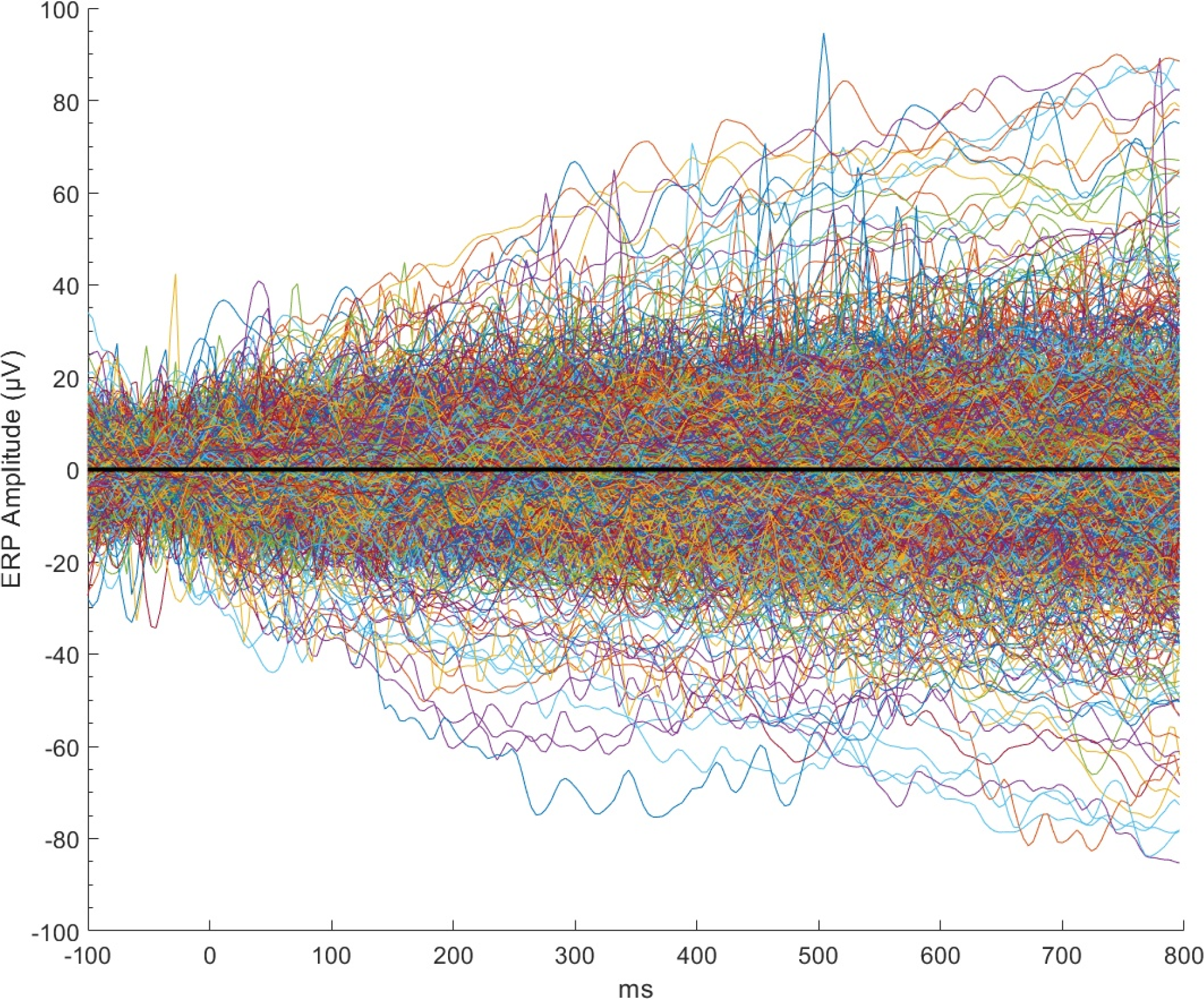
ERP traces (N=12,520) after plus-minus averaging reflect noise (colors). A black line reflecting average noise is overlaid. The average of the noise was approximately zero, indicating that the signal was indeed subtracted out by the plus-minus averaging procedure and that only noise remained in the data.

**Figure 4:**
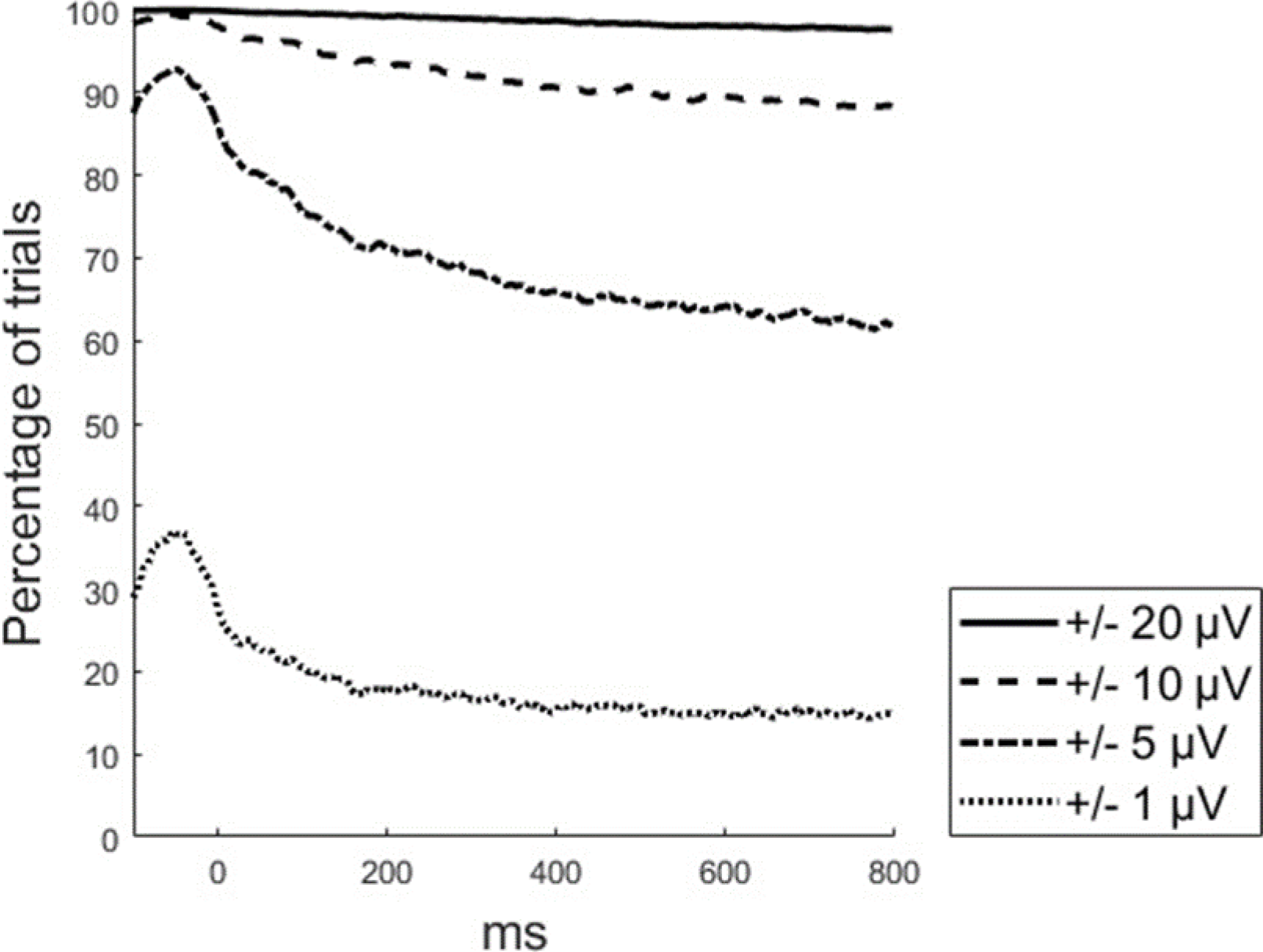
Line plot showing the percentage of experimental trials within the voltage bins of ±20 µV, ±10 µV, ±5 µV, and ±1 µV. Approximately only 2% of trials exceeded the ±20 µV threshold at any given point.

### 2.6. Monte Carlo simulation

To simulate separate ERP experiments using either within-subject or between-subjects designs, we randomly sampled subsets of subjects from the larger data set described above. For each subject we randomly sampled subsets of trials to which we added known effects to simulate LPP responses to different conditions, such as neutral and emotional stimuli. Each simulated experiment included a specific effect size, number of trials, and number of subjects. For within-subject designs, we selected twice the number of trials from each randomly sampled subject in order to simulate two experimental conditions. For the between-subjects simulations, we sampled twice the number of subjects to simulate two experimental groups. For the within-subject analysis, we modeled effect size by adding one-half the simulated effect size to the LPP of one condition and by subtracting one-half of the simulated effect size from the other.

Similarly, between-subjects effect sizes were modeled by adding one-half the simulated effect size to the LPP of one group and subtracting one-half the simulated effect size from the LPP of the second group. The size of the artificial effects ranged from 0 to 3 µV, in increments of 0.1 µV; the number of trials ranged from 5 to 40 trials per condition, in increments of five trials; and the number of subjects in each experiment ranged from 10 to 100, in 10-subject increments from 10 to 50 and a further increment of 50 to reach 100. Combining all parameters led to a total of 1,488 simulated experiments. Each simulated experiment was repeated 1,000 times. All Monte Carlo simulations were performed using MATLAB R2018b (The MathWorks, Inc., Natic, MA, USA).

### 2.7. Statistical analysis

For each simulated experiment, we tested for statistically significant effects (p<.05) using one-way repeated measures ANOVA for experiments simulating within-subject effects and one-way ANOVA for experiments simulating between-subjects effects. It is important to note that these between-subjects ANOVAs essentially model the interaction effect of groups by condition with two groups and two conditions. The effects added to the trials of the participants in each group can be thought of as the voltage difference between images belonging to two different categories. This procedure allowed us to estimate the probability of obtaining a statistically significant outcome (α = 0.05) for each combination of parameters. All ANOVAs were performed in R 3.5.0 (R Core Team, 2018). For each experimental condition, the percentage of F values at or above the critical F value was calculated to represent statistical power.

## 3. Results

### 3.1. Within-subject analyses

As shown in Figure 5, within-subject analyses revealed that when only 10 subjects were included in the experiment, 80% power was only achieved for differences in effect sizes larger than 1 µV, even when a large number of trials (40) was included for each experimental condition. With smaller numbers of trials and smaller differences in effect sizes, sufficient power to detect the differences could not be achieved.

**Figure 5:**
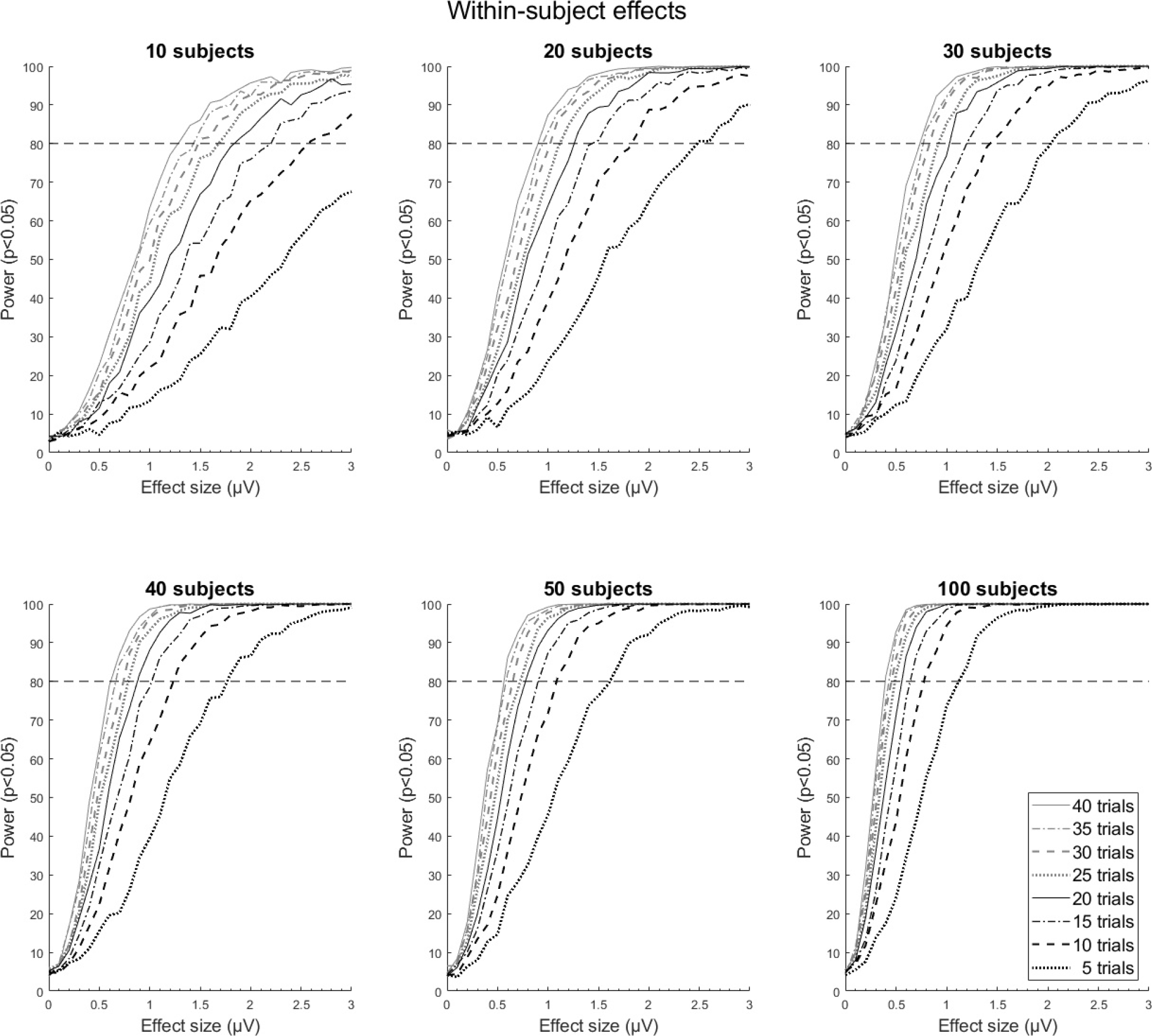
Within-subject analysis of statistical power by effect size at 5-40 trials per condition and for 10-100 subjects.

We found that, as the number of subjects increased, statistical power reached an asymptote at 100% at 1.5 µV for the experiments with a higher number of trials. This asymptote became apparent for experiments with smaller numbers of trials as the number of subjects increased and became evident for effect sizes as small as 1 µV with greater numbers of trials.

### 3.2. Between-subjects analyses

Overall, statistical significance is harder to achieve in between-subjects experimental designs than within-subject designs, and this was reflected in our results. Figure 6 shows that statistical power was only achieved at higher effect sizes, greater numbers of trials, and relatively large sample sizes compared with what was observed in our within-subject analyses. The slopes shown in Figure 6 were generally less steep than the slopes shown in Figure 5, indicating that an increase in the size of the difference between conditions did not affect statistical power as dramatically in between-subjects designs as it did in within-subject designs. However, the overall trend of slopes increasing and shifting to the left with increasing sample sizes was conserved. At lower numbers of subjects per group, 80% statistical power was much harder to achieve between-subjects than within-subjects and was only achieved with greater numbers of trials. Starting at 40 subjects per group, statistical power reached an asymptote of 100% at 20 or more trials for effect sizes greater than 1.5 µV. This asymptote shifted to include smaller effect sizes and lower numbers of trials as subjects were added to the experiment, with 100 subjects per group reaching an asymptote at effect sizes as low as 1 µV. At 100 subjects per group, slopes were steepest, with 80% power achieved at effect sizes as low as 0.5 µV for experiments with 40 trials per condition.

**Figure 6:**
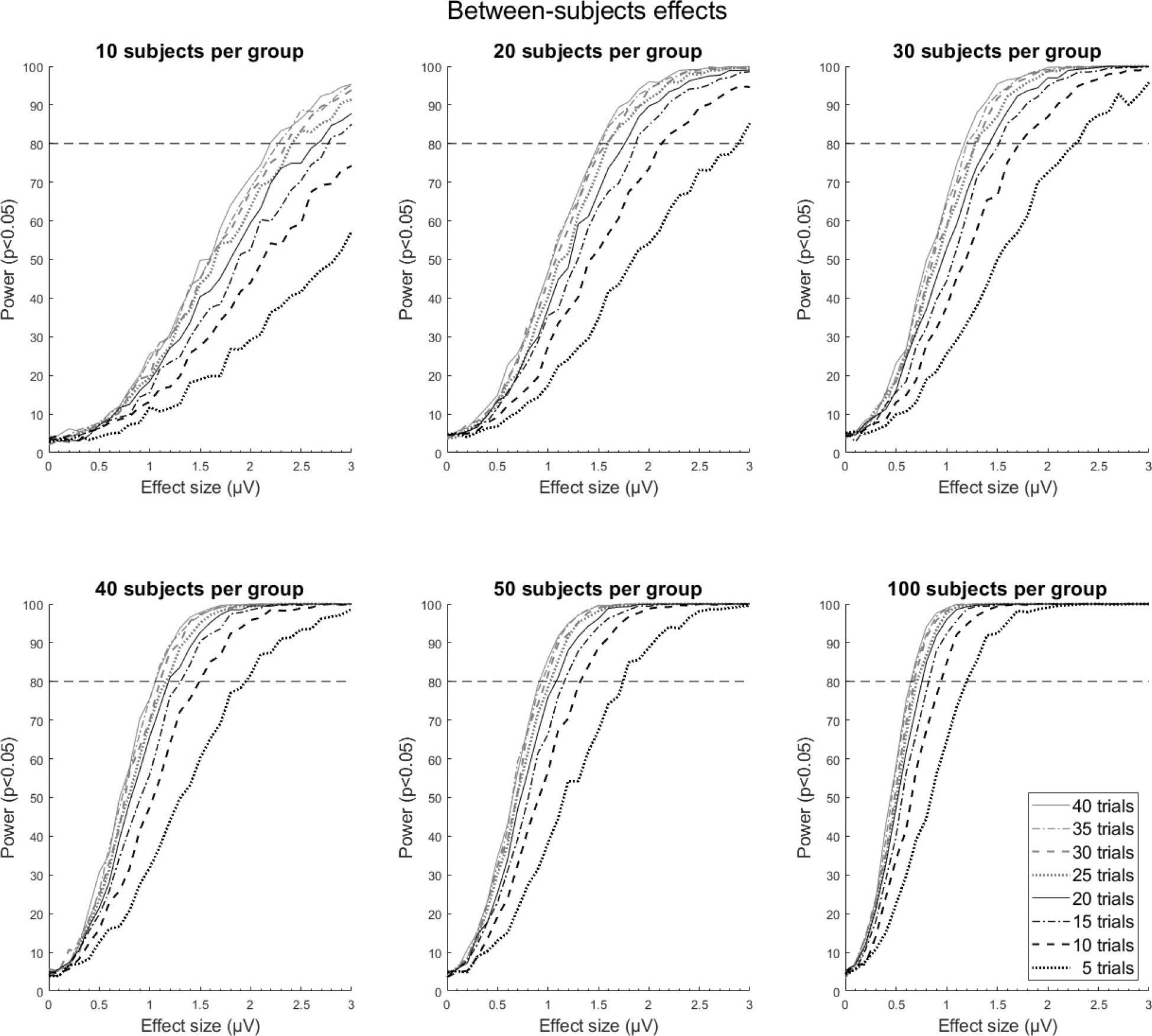
Between-subjects analysis of statistical power by effect size at 5-40 trials and 10-100 subjects per group

### 3.3. Summary

Predictably, our results showed that increasing the number of trials and subjects increased statistical power and that statistical power was greater for larger effect sizes. Also, as expected, statistical power of at least 80% could be achieved at lower effect sizes, population sizes, and trial numbers in within-subject as compared with between-subjects experiments. Furthermore, we found that, in both within-subject and between-subjects experiments, an increase in subjects more rapidly increased the statistical power at the lower range of effect sizes (<1 µV) until the power later reached an asymptote at the higher range of effect sizes (>1.5 µV).

## 4. Discussion

The present study was concerned with how best to optimize the parameters that affect statistical power in ERP experiments that use the amplitude of the LPP to assess affective processes in within-subject and between-subjects designs. We adopted the simulation approach recently used by Boudewyn and co-workers (2018), but we investigated the LPP, an ERP component that is heavily studied in the domain of affective neuroscience and, compared with the ERN, has more modest effect sizes.

Detecting differences in the amplitude of the LPP evoked by emotional versus neutral stimuli between groups often means investigating modest differences; accordingly, achieving sufficient statistical power under such conditions requires increasing the number of subjects and/or including a greater number of trials in the experiment. Here we provide a useful tool that researchers can use to evaluate the tradeoffs and achieve their research objectives.

### 4.1. Different ERP components have different dynamics with respect to power

Previous studies have investigated statistical power for the ERN and LRP components (Boudewyn et al., 2018), and it is interesting to note that, in the Boudewyn et al. study, the slope of the relationship between the statistical power and the number of trials was steepest at the middle range of the effect sizes (3-5 µV) and numbers of trials (10-12) investigated. In contrast, the present study found steeper slopes at the higher range of the numbers of subjects (50+) and trials (20+) but at lower effect sizes (0.4-1 µV). Therefore, the dynamics of study parameters, as they relate to statistical power, are different among ERP components. It is likely that these dynamics are related both to the properties of the ERP components themselves and to the noise present in the data. As such, experiments that assess the LPP may benefit more from increasing numbers of trials and subjects than would experiments that focus on larger amplitude components such as the ERN. Thus, the signal-to-noise ratio (SNR) for the LPP may increase more with increased numbers of trials and subjects as the variability between individuals and the noise within the component itself are averaged out.

### 4.2. Affective neuroscience

Our results provide guidelines that neuroscientists can employ when designing experiments that use the LPP to investigate affective processes or when evaluating the results of these experiments. Our findings indicate that between-subjects comparisons that include, say, 10 subjects per group are extremely unlikely to produce meaningful results. As pointed out by Ioannidis et al. (Ioannidis, 2005), when experiments are grossly underpowered, a statistically significant result is likely to be artifactual. Even with 20 or 30 subjects per group, sufficient statistical power can be achieved only for effect sizes larger than 1 µV, an effect larger than the LPP difference that, in our data, separated low arousing from neutral stimuli (Figure 2). For investigators studying more modest differences, such as those often observed in interaction effects in both within-and between-subjects designs), our results indicate that, at a minimum, 40 trials and 50 subjects per group are needed to achieve sufficient statistical power.

Our findings related to the SNR of the LPP signal also inform the field of affective neuroscience regarding the trade-off between adding subjects vs adding trials to an experiment. As shown in Figure 6, 5 trials per condition and 10 subjects per group achieves a statistical power of approximately 25% at an effect size of 2 microvolts. However, an experiment with the same parameters that includes 20 subjects per group achieves nearly 55% statistical power: by doubling the population size, the power increases approximately twofold in this example. Meanwhile, if the number of trials per condition is doubled to 10 trials per condition while still using 10 subjects per group and looking at a 2 microvolt effect size, statistical power is approximately 40%. Thus, our results indicate that adding subjects to an experiment has a greater effect on the statistical power than increasing the number of trials would. However, due to the difficulties that come with the recruitment of human subjects, doubling the sample size may not be feasible or effective, and some labs may favor adding trials to their experiments instead. Conversely, depending on the number of conditions used in the experiment, adding trials may not be feasible, as this could excessively increase the duration of the experiment. The results that we presented here offer researchers the opportunity to more precisely estimate the impact that decisions about important parameters in an experiment have on statistical power.

One objective of this study was to simulate data with a higher degree of noise and with modest effect sizes, as many investigators may be interested in how to sufficiently power studies investigating small LPP amplitude differences or may work with noisy data. Hence, our results might be less informative when very robust effects (e.g., those greater than 3 microvolts) are under investigation or when noise in the data is minimal. Furthermore, it is likely that different ERP components might show different dynamics with respect to power, and as such future studies should specifically investigate statistical power for more components.

## 5. Conclusions

By sufficiently powering clinical affective neuroscience studies, investigators will collect more reliable results, thereby improving the reproducibility of their research findings. Our findings may help researchers planning and evaluating the results of experiments that use the LPP as an index of motivational relevance and may ultimately foster the translation of results from basic science experiments to evidence-based treatments of disorders characterized by altered affective processing. Careful consideration of statistical power furthers the ultimate goal of translational affective neuroscience research: to bolster innovation in the psychiatric domain.

## Acknowledgements

Department of Scientific Publications, The University of Texas MD Anderson Cancer Center

